# Neural control of respiration in *Drosophila*

**DOI:** 10.64898/2026.07.28.740380

**Authors:** Yichen Luo, Anne Sustar, Juan Ispizua, Gayathri Kondakath, Yu Yang, Floris van Breugel, John C. Tuthill

**Affiliations:** Department of Neurobiology and Biophysics, University of Washington, Seattle, WA 98195, USA; Neural Systems & Behavior Course, Marine Biological Laboratory, 7 MBL Street, Woods Hole, MA 02543, USA; Department of Biology, Tufts University, Medford, MA 02155, USA; Department of Mechanical Engineering, Johns Hopkins University, 3400 North Charles Street, MD 21218, USA; Department of Mechanical Engineering, University of Nevada Reno, Reno, NV 89512, USA

## Abstract

Adaptive control of breathing is essential for life, yet the neural circuits that couple respiration to locomotion and internal physiological state remain poorly understood. Insects regulate gas exchange through spiracles: valve-like openings in the cuticle that allow oxygen uptake while minimizing water loss. We find that spiracle opening in *Drosophila* is dynamically matched to flight power and reduced by dehydration. We identify the motor neurons that innervate the spiracle muscles and show that their optogenetic activation closes all eighteen spiracles and rapidly limits flight power. Inhibitory interneurons transmit descending flight commands to the spiracle motor neurons, opening spiracles in proportion to metabolic demand. A parallel interoceptive pathway converges on the same interneurons to close the spiracles and suppress flight. Feedforward descending commands thus couple flight power to spiracle opening, while interoceptive feedback pathways close spiracles to balance oxygen supply against water loss. This compact circuit architecture may reflect a general solution for matching respiration to the competing demands of locomotion and water conservation.

## Main

Breathing is among the most fundamental and evolutionarily conserved of all motor behaviors (*1*). In all terrestrial animals, motor control of respiration is tightly controlled by the nervous system, which adjusts respiratory behavior based on metabolism (*2–4*), arousal (*5*), and exertion (*6*, *7*). In vertebrates, the respiratory rhythm is generated by brainstem circuits (*8*) that adjust breathing rate to match locomotor state (*6*, *9*) as well as oxygen and carbon dioxide levels in the blood (*10*). This tight coupling between respiration, metabolism, and locomotion allows mammals, including humans, to adjust their oxygen consumption by 10-20-fold between rest and peak exertion (*11*, *12*). Feedback mechanisms have been characterized in detail, but feedforward control has proven far more elusive (*13*). Feedback alone also cannot account for the tight coupling among respiration, metabolism, and locomotion, since a controller that waits for internal gases to change will always lag demand.

While larger terrestrial vertebrates rely on circulation of oxygen in the blood, insects directly absorb oxygen into their internal tissues through extensive networks of air-filled tubes called trachea (*14–16*). Although highly efficient, direct gas exchange exposes a large internal surface area to the outside world, which increases water loss (*17*, *18*). To regulate air flow into the tracheal system, insects possess small valve-like openings on the cuticle called spiracles. Classic studies showed that spiracles can be opened and closed depending on internal hydration state (*17*, *18*) and the motor exertion (e.g., flight, walking, quiescence) of the animal (*4*, *19*, *20*). This control is particularly critical during flight, one of the most metabolically expensive behaviors in the animal kingdom (*19–21*). Winged insects consume ∼5 mL O₂/g/hr at rest (*22*, *23*), comparable to the peak oxygen consumption of an elite human athlete. They can also increase VO₂ more than tenfold during flight — among the largest metabolic dynamic ranges in the animal kingdom (*24*). Electrophysiological recordings from larger insects have identified candidate sites of respiratory-locomotor coupling (*25*), but without genetic access to these neurons, their causal roles could not be established.

*Drosophila melanogaster*, with its compact nervous system, comprehensive neural wiring diagrams (*26–33*), and powerful molecular genetic toolkit (*34–38*), is uniquely positioned to fill this gap. In the wild, *Drosophila* can disperse remarkably long distances, traveling up to ∼12 km in a single flight across dry desert terrain (*39*). Sustaining flight under these conditions requires balancing the high oxygen demand of the flight muscles against the risk of water loss through the spiracles. This tradeoff is likely to become increasingly critical as insects face rising temperatures and aridity due to climate change (*40*).

Fruit flies breathe through nine pairs of spiracles distributed along the body (**Fig. 1A**). The two largest spiracles are located on the thorax: the mesothoracic spiracle (Sp1), positioned dorsal to the foreleg, and the metathoracic spiracle (Sp2), beneath the haltere (**Fig. 1B**). Together, the two pairs of large thoracic spiracles account for ∼95% of the total spiracular surface area, while the seven pairs of smaller abdominal spiracles comprise the remaining 5% (*4*). Thoracic spiracles are covered by filter hairs called trichomes and guarded by two membranous flaps that contain resilin (**Fig. 1C, top**), an elastic protein that makes cuticle more deformable (*41*, *42*). Both flaps are mechanically linked to a small muscle located ventral to the spiracle (**Fig. 1C, bottom**), which is presumed to regulate airflow by actively opening or closing the flaps. Prior work used respirometry to measure CO_2_ and H_2_O flux in flying *Drosophila*, suggesting that flies open their spiracles to increase respiration during flight (*4*, *20*, *43*, *44*). However, the dynamics of spiracle opening and closing have not previously been measured in *Drosophila*, and the neural circuits that control fly respiration remain unknown.

**Fig. 1.**
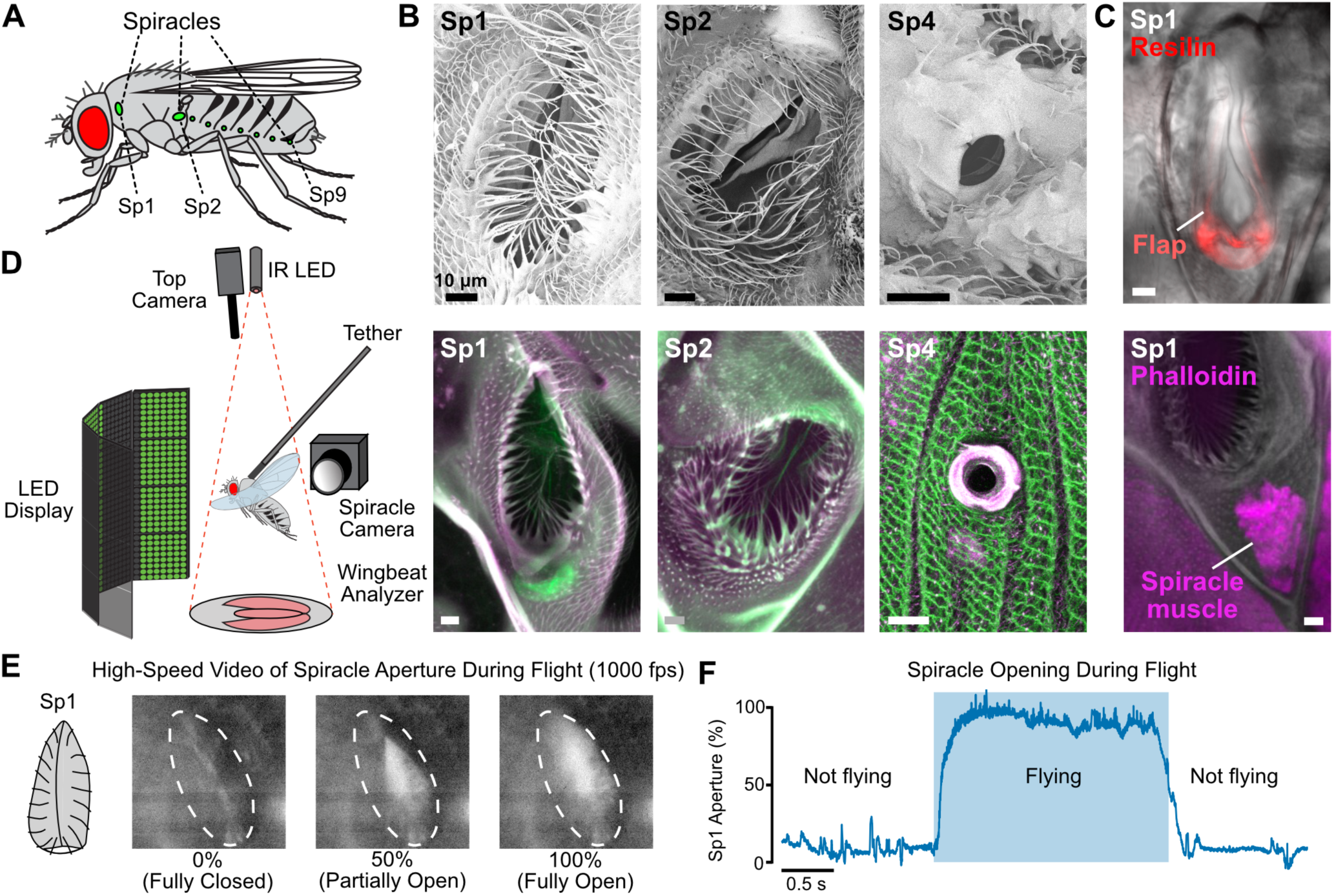
The spiracles of adult *Drosophila***. (A)** The two thoracic spiracles, Sp1 and Sp2, comprise ∼95% of total spiracular surface area. **(B)** Sp1, Sp2, and an abdominal spiracle (Sp4), imaged by scanning electron microscopy (SEM, top) and confocal microscopy (autofluorescence, bottom). **(C)** The Sp1 closer muscle and resilin-rich flaps indicate a deformable, spring-loaded valve. **(D)** Closed-loop visual flight arena: wingbeat amplitude, frequency, and Sp1 aperture are recorded simultaneously. **(E)** Sp1 aperture, filmed at wingbeat frequency, is quantified as mean pixel intensity within an ROI around the orifice. **(F)** Sp1 opens at flight onset, stays open throughout the bout, and closes at flight cessation. Results were similar with Sp2 (**Movie S1**).

### Spiracle opening is modulated by internal physiological state and flight demand

We used high-speed video (**Fig. 1D**) to record the thoracic spiracles (Sp1, Sp2; **Fig. 1E, Fig. S1A**) in tethered flies. When the fly was at rest, its spiracles remained mostly, though not completely, closed (**Fig. 1F**). Thoracic spiracles opened at flight initiation and closed shortly after flight cessation (**Fig. 1F, Movie S1, S2**), similar to what has been observed in larger insects (*21*, *45–47*). During longer periods of sustained flight, Sp1 aperture fluctuated continuously even when wingbeat amplitude (WBA) and frequency (WBF) were stable (**Fig. S1B**), showing that spiracle aperture is actively modulated during flight.

To test whether the spiracles track flight demand in real time, we presented tethered flying flies with a vertically oscillating visual stimulus (**Fig. 2A**) that drives an optomotor lift response: upward visual motion causes flies to bilaterally increase their WBA, raising flight power to generate lift (*48*). The mesothoracic spiracle Sp1 closely tracked these changes, opening when flight power was high and closing when flight power was low (**Fig. 2B, Fig. S1C, Movie S3**). Across flies, Sp1 aperture, WBA, and flight power were all strongly and consistently correlated with stimulus velocity (**Fig. 2C**), whereas the correlation with WBF was weaker and more variable. Sp1 aperture lagged the visual stimulus by a delay comparable to that of WBA and flight power (**Fig. 2D**), rather than trailing behind them. The metathoracic spiracle Sp2 was similarly modulated (**Movie S4**). Together, these results show that thoracic spiracle aperture is continuously matched to flight power on a moment-to-moment timescale, with a latency consistent with feedforward coordination of respiration and flight rather than reactive metabolic feedback.

**Fig. 2.**
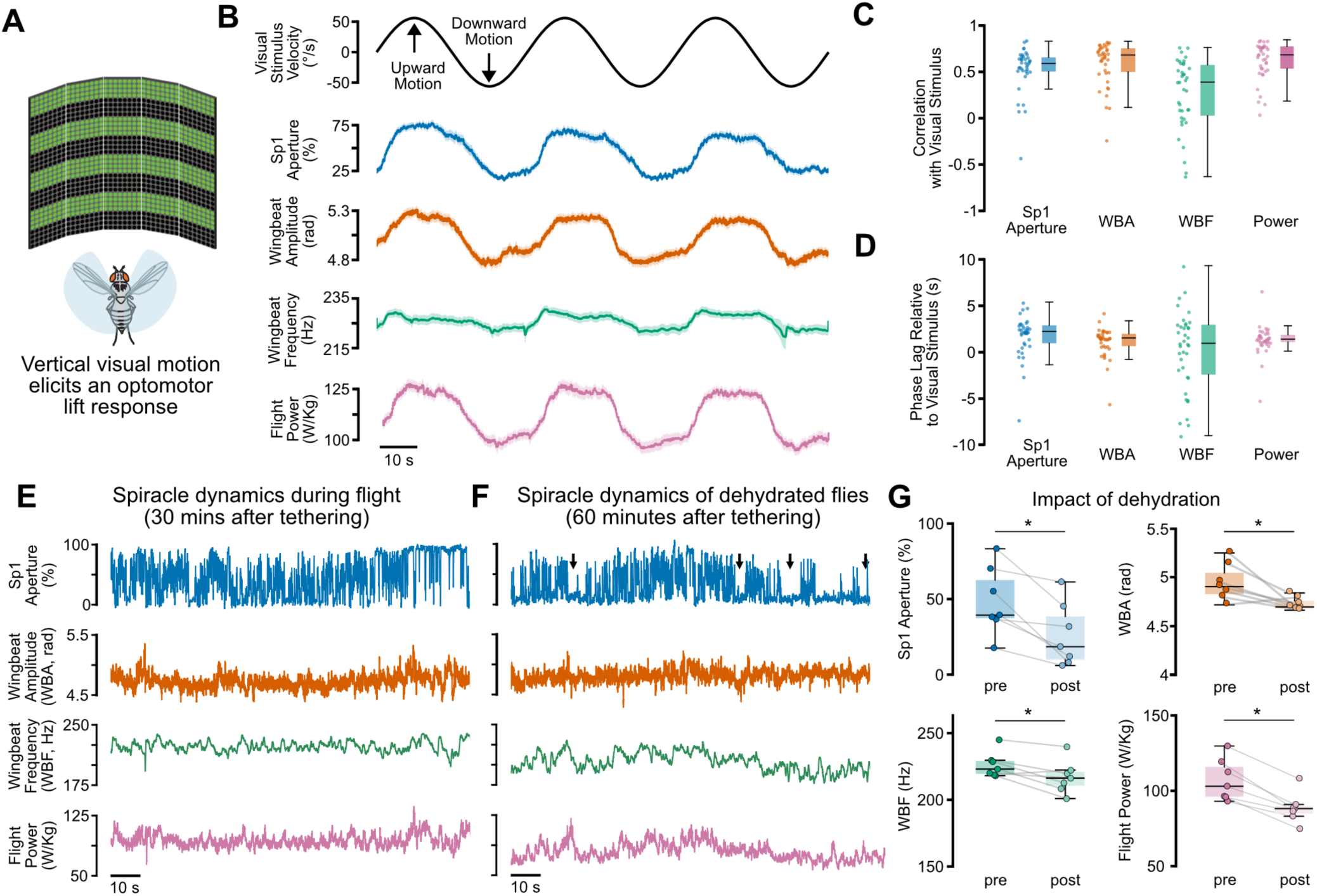
Thoracic spiracle aperture is regulated on slow and fast timescales**. (A)** Optomotor stimulus that modulates flight power: vertically moving bars at sinusoidal velocity, with simultaneous closed-loop stripe fixation. **(B)** Sp1 aperture covaries with stimulus velocity, WBA, and flight power. n = 38 flies. **(C)** Correlations with Sp1 aperture, WBA, and flight power are high and consistent; WBF correlates more weakly and variably. n = 38. **(D)** Sp1 aperture lags the stimulus by a delay comparable to WBA and flight power. n = 38. **(E)** At ∼30 min after tethering, Sp1 fluctuates between fully open and closed, with fast (phasic) flutter and slower (tonic) modulation, while wing kinematics remain stable. (**F**) After 30 mins of desiccation, mean aperture is reduced and sustained closures occur (arrows) despite stable WBA and WBF. **(G)** Sp1 aperture, WBA, WBF, and flight power all decline significantly after desiccation. n = 7; paired Wilcoxon test, *p < 0.05.

We next asked whether spiracle opening is also shaped by internal state, focusing on hydration because of the tradeoff between oxygen uptake and water conservation. We recorded the Sp1 spiracle of tethered flies before and after dehydrating them in a desiccation chamber for 30 minutes. After the dehydration treatment, spiracle aperture was reduced and the spiracle closed for prolonged periods (**Fig. 2E-F, arrows; Movie S5**). This decline in Sp1 opening was accompanied by significant reductions in WBA, WBF, and flight power (**Fig. 2G**). Thus, dehydration suppresses spiracle opening and flight together, indicating that spiracle aperture is shaped not only by moment-to-moment flight demand but also by a slower, hydration-dependent signal.

### Identification of motor neurons that close the spiracles

As an entry point to the neural circuits that control respiration, we sought to identify the motor neurons that innervate the spiracle muscles. We used a connectome of the male *Drosophila* ventral nerve cord (VNC) (*32*, *49*) to search for motor neurons that projected efferent axons through peripheral nerves (**Fig. 3A**). Focusing on axons that project through the wing and haltere nerves resulted in ∼40 bilateral pairs, of which 35 were previously annotated as wing or haltere motor neurons. Among the five remaining unannotated pairs, two shared highly similar presynaptic input patterns (cosine similarity; **Fig. S2A**), suggesting a common function. Applying the same common input criterion to the abdominal nerves identified seven additional bilateral candidate spiracle motor neuron (SpMN) pairs, matching the seven abdominal spiracles. Imaging a driver line that labels all motor neurons (GluRIIB-GAL4(*50*)) confirmed that each spiracle muscle is innervated by a single axon (**Movie S6**). In addition to the glutamatergic motor neurons, it is also possible that spiracle muscles are innervated by efferent aminergic neurons, as in the locust (*51*).

**Fig. 3.**
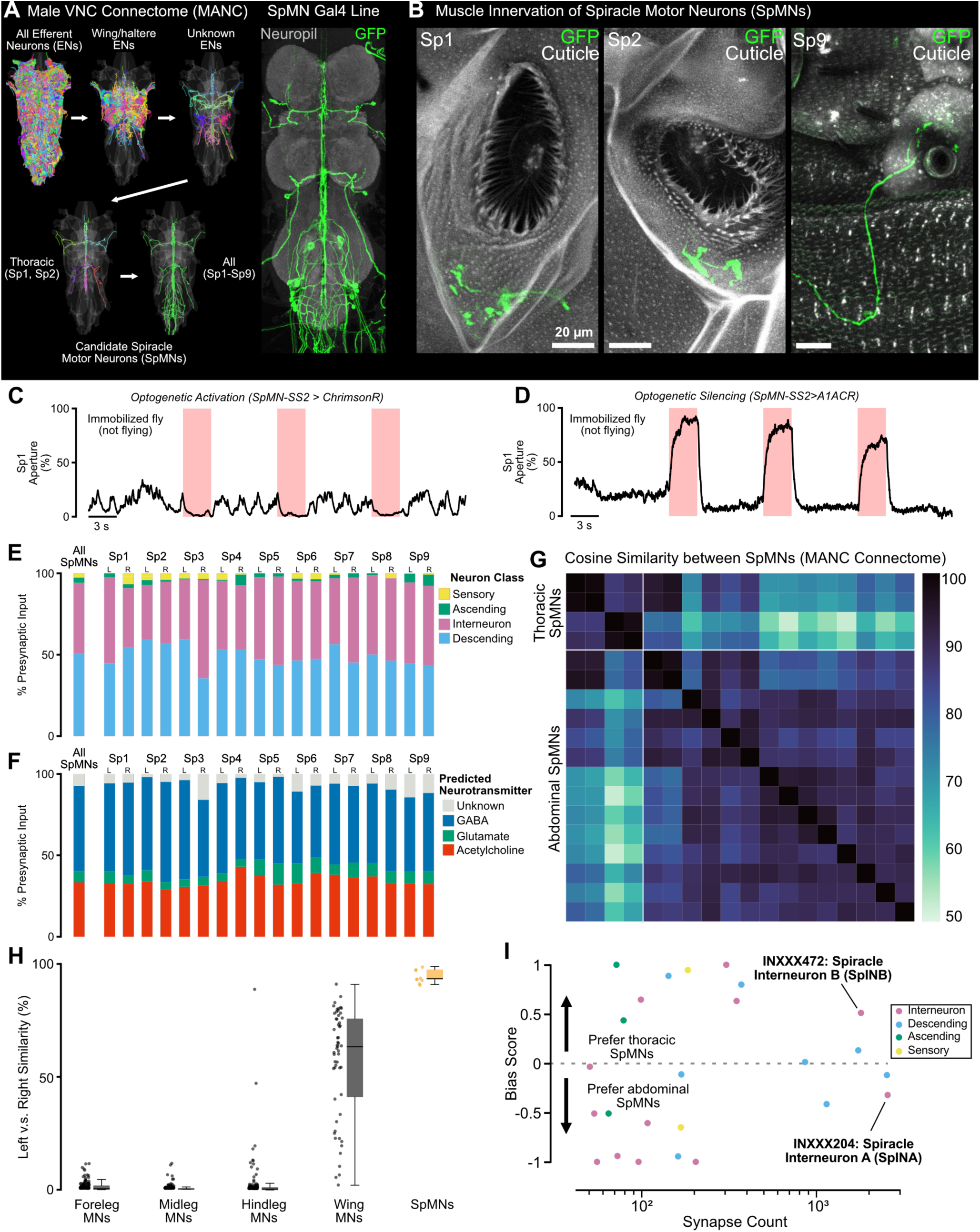
Connectome-guided identification of spiracle motor neurons and their presynaptic connectivity patterns. **(A)** A connectome screen of MANC narrowed ∼800 VNC efferents to ∼40 bilateral pairs, then used presynaptic input similarity to identify 2 thoracic and 7 abdominal SpMN pairs. We generated a Split-Gal4 line (SpMN-SS1-GAL4) that labels all SpMNs. **(B)** A second split line (SpMN-SS2-GAL4) labels axons innervating Sp1, Sp2, and an abdominal spiracle muscle, confirming SpMN identity. **(C)** Optogenetic activation of SpMNs closes Sp1. (**D**) Silencing opens Sp1. **(E)** SpMNs receive ∼40% of presynaptic input each from descending neurons and VNC interneurons. **(F)** SpMNs receive predominantly cholinergic (∼30%) and GABAergic (∼50%) input. **(G)** Presynaptic input vectors cluster into thoracic and abdominal blocks, indicating differential upstream control, though overall similarity across SpMNs is high. **(H)** Bilateral SpMN pairs show the highest left–right input similarity of any motor neuron class examined, suggesting tightly coordinated bilateral spiracle control. **(I)** SpINA (GABAergic) preferentially targets abdominal SpMNs; SpINB (GABAergic) preferentially targets thoracic SpMNs. Both provide strong input to all SpMNs.

Once we identified candidate SpMNs in the connectome, we used NeuronBridge (*37*) to match these cells to a library of genetic driver lines, from which we generated two Split-Gal4 lines that specifically label SpMNs(SpMN-SS1, SpMN-SS2; **Fig. 3A**, right; **Fig. S2B**). GFP expression confirmed that the axons of these motor neurons innervate thoracic and abdominal spiracles (**Fig. 3B**). Optogenetic activation of SpMNs using the red-shifted channelrhodopsin ChrimsonR (*52*) closed thoracic and abdominal spiracles (**Fig. 3C, Movie S7**), and silencing SpMNs using the red-shifted anion channelrhodopsin A1ACR (*53*) opened spiracles to their maximum extent (**Fig. 3D**). These results indicate that *Drosophila* spiracles are controlled by closer muscles.

We used the male VNC and full CNS (*54*) connectomes to investigate the organization of presynaptic neurons that synapse on SpMNs. All 18 SpMNs received similar fractions of input from descending neurons, interneurons, ascending neurons, and sensory neurons, with descending neurons and intrinsic VNC interneurons each contributing roughly 40% of all synaptic input (**Fig. 3E, Fig. S2C-D**). Grouping presynaptic input by predicted neurotransmitter revealed ∼30% cholinergic and ∼50% GABAergic input (**Fig. 3F, Fig. S2E**). At the population level, the similarity of presynaptic inputs (**Fig. 3G)** was substantially higher for bilateral left-right SpMN pairs than for bilateral pairs of leg or wing motor neurons (**Fig. 3H**). This pattern of connectivity to SpMNs is consistent with coordinated control of bilateral spiracle pairs; in other words, our finding that SpMNs receive input from the same presynaptic cells suggests that left/right pairs of spiracles open and close together.

Despite the overall similarity of presynaptic input to spiracles, connectivity was more similar within thoracic SpMNs and abdominal SpMNs than across the two subclasses (**Fig. 3G, Fig. S2F**). This suggests that while all SpMNs share a high degree of common input, thoracic and abdominal spiracles may be differentially controlled under certain conditions. Thoracic spiracles deliver air to the flight muscles and head, while abdominal spiracles supply the abdomen, so differential control could allow flies to regulate airflow in different body segments to meet different physiological needs (e.g., thinking versus digestion). To identify neurons that could mediate differential control of thoracic and abdominal spiracles, we computed a bias score reflecting the fraction of each presynaptic neuron’s output directed to thoracic versus abdominal SpMNs (**Fig. 3I**). Two GABAergic interneurons stood out with the highest bias scores, synapse counts, and fractions of total output directed to SpMNs: we call these cells the spiracle interneurons, SpINA (INXXX204) and SpINB (INXXX472). SpINA preferentially targets abdominal SpMNs, while SpINB targets thoracic SpMNs. Several additional interneurons showed bias between Sp1 and Sp2 (**Fig. S2G**), suggesting that even the two thoracic spiracles may be differentially controlled.

### Spiracle control and motor behavior

Having identified the SpMNs and characterized their presynaptic connectivity, we asked whether spiracle opening is required to sustain flight power. We used optogenetics to activate and silence SpMNs during tethered flight, restricting illumination to the VNC to target the SpMNs while sparing off-target cells in the brain (**Fig. 4A**). We used our two Split-Gal4 driver lines, which differ in their expression patterns: SpMN-SS1 labels all SpMNs throughout the body, whereas SpMN-SS2 labels the thoracic SpMNs but stochastically omits ∼4-6 abdominal SpMNs (**Fig. 4C, Fig. S2B**). This difference allowed us to compare the consequences of complete versus partial spiracle closure.

**Fig. 4.**
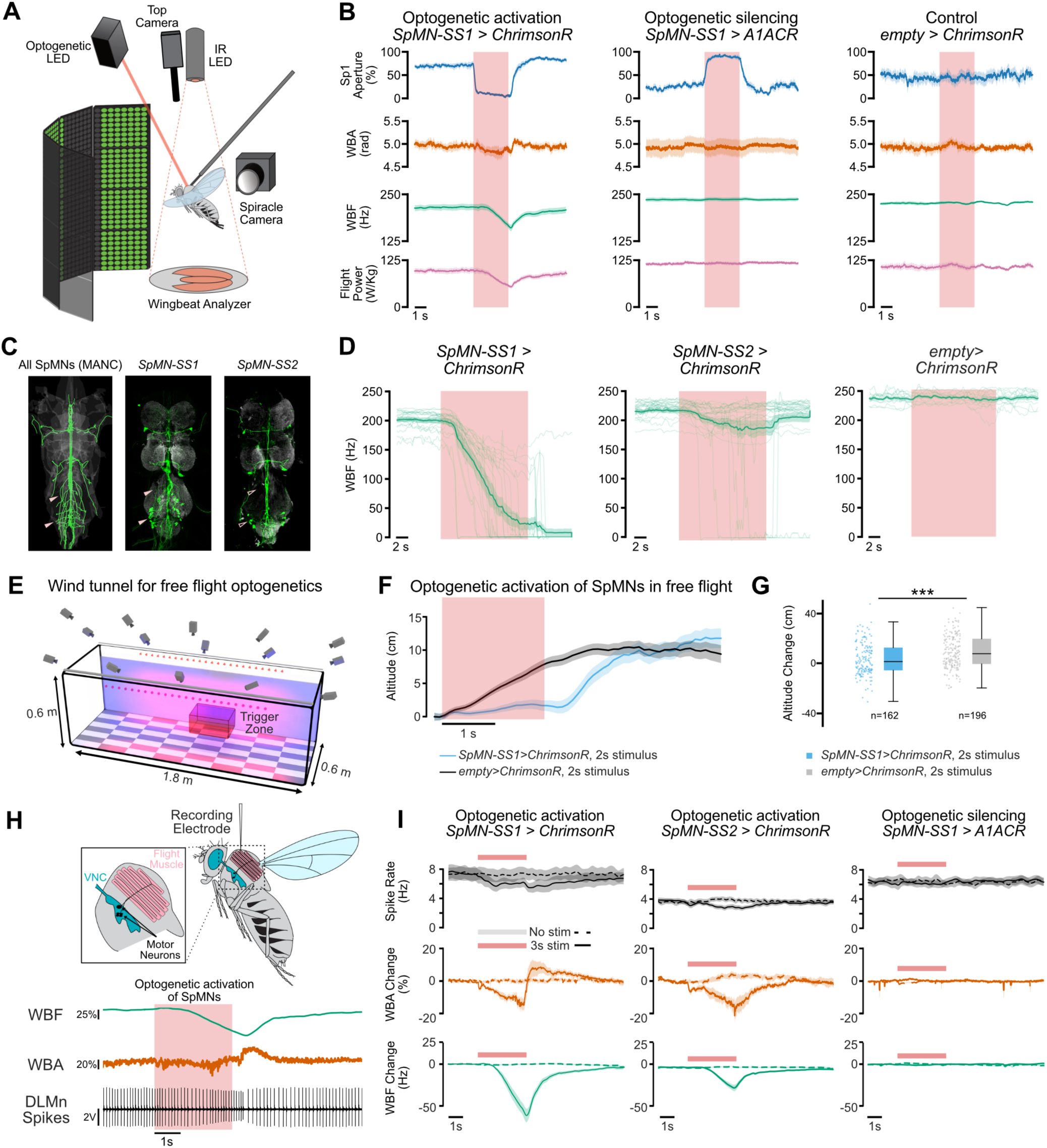
Spiracle closure rapidly reduces flight power and can uncouple motor neuron spiking from aerodynamic output. **(A)** Optogenetic activation/silencing during flight. **(B)** SpMN closure rapidly decreases WBA, WBF, and flight power; opening and controls are unaffected. n = 10–11. **(C)** SpMN-SS2 labels fewer SpMNs than SpMN-SS1, stochastically omitting several abdominal motor neurons (arrows). **(D)** Activating all SpMNs (SpMN-SS1) typically stops flight within 10 s; activating a thoracic-biased subset (SpMN-SS2) has a weaker effect. n = 22 for *SpMN-SS1>ChrimsonR*, n = 24 for *SpMN-SS2 > ChrimsonR*, n = 10 for *empty > ChrimsonR*. **(E and F)** In free flight, optogenetic spiracle closure causes a pronounced loss of altitude relative to controls. **(G)** Closing spiracles significantly reduces flight altitude relative to controls. Welch’s t-test, ***p < 0.001. **(H and I)** During spiracle closure, DLMn spike rate falls modestly and recovers slowly, remaining below baseline after stimulus offset, while WBA and WBF drop sharply. Spike rate is unchanged when spiracles are opened instead. n = 11 for *SpMN-SS1>ChrimsonR*, n = 15 for *SpMN-SS2 > ChrimsonR*, n = 13 for *SpMN-SS1 > A1ACR*.

When we closed all the spiracles through optogenetic activation of SpMN-SS1, wingbeat frequency and amplitude began to decrease within < 2 s. Both WBA and WBF recovered after stimulus offset, with amplitude recovering faster than frequency (**Fig. 4B, Movie S8**). Prolonged (10 s) activation of SpMN-SS1 typically terminated flight altogether (**Fig. 4D, left**). In contrast, activation of SpMN-SS2, which spares a subset of abdominal spiracles, reduced WBF but rarely stopped flight (**Fig. 4D, right**). This result suggests that residual gas exchange through the remaining open abdominal spiracles can sustain flight even when the thoracic spiracles are fully closed (*43*). Silencing SpMNs during flight fully opened the spiracles but had no detectable effect on WBF or WBA (**Fig. 4B, middle, Movie S8**), and control flies did not change spiracle aperture or WBA/WBF in response to the red-light stimulus. Because wingstroke kinematics and flight behavior differ between tethered and free flight, we repeated the activation experiments in a free-flight arena (**Fig. 4E**). SpMN-SS1 activation caused flies to lose altitude relative to controls (**Fig. 4F–G**), establishing that spiracle closure compromises flight performance in both tethered and free flight.

A decrease in wingbeat frequency and amplitude could result from decreased excitatory drive to the flight muscles, reduced muscle contractility, or both. We sought to distinguish between these alternatives using extracellular recordings from the dorsal longitudinal muscle (DLM) motor neurons (DLMns), which drive the indirect flight muscles (**Fig. 4H**). In response to both SpMN-SS1 and SpMN-SS2 activation, spiracle closure produced a modest but prolonged decrease in DLMn firing rate that outlasted the optogenetic stimulus (**Fig. 4I, Fig. S3**), whereas silencing left firing rate unchanged. The decrease in DLMn spike rate was too small to account for the much larger decline in WBF and flight power. Notably, while spike rate fell on average, individual flies showed both increases and decreases, yet flight power declined in all cases (**Fig. S3A**). Indeed, the relationship between DLMn firing and aerodynamic output was changed during and after closure (**Fig. S3B**). For a given firing rate, flies produced substantially less power during and after the optogenetic stimulus than before it (**Fig. S3C**). These results indicate a failure of force production downstream of motor neuron drive. In other words, our data suggest that spiracle closure impairs flight through a peripheral failure of flight muscle power generation, consistent with oxygen limitation of flight muscles.

### A feedforward circuit that matches spiracle aperture to expected metabolic demand

Having found that thoracic spiracle aperture is continuously matched to flight power and that closing spiracles impairs flight, we asked how this coupling is implemented at the circuit level. Our connectome analysis identified the left/right pair of SpINB neurons as a top input to the thoracic SpMNs (**Fig. 5A**). The SpINB cells are strongly connected to SpMNs: ∼80% of their output synapses is onto SpMNs, and the bias score indicates that SpINB preferentially synapse with thoracic SpMNs, which are likely involved in regulating respiration during flight. The SpINB cells receive direct synaptic input from DNg02, a class of descending neurons previously shown to control flight power (*55*, *56*). The DNg02 neurons also synapse directly onto the indirect flight motor neurons (DLMns, **Fig. 5B**). Finally, the SpINB cells receive input from other descending neurons, DNg27 and DNge137, which synapse strongly with SpMNs (**Fig. S2C**). Overall, SpINB is well positioned to relay a feedforward copy of the descending flight command to set thoracic spiracle opening in proportion to commanded flight power (**Fig. 5A-C)**.

**Fig. 5.**
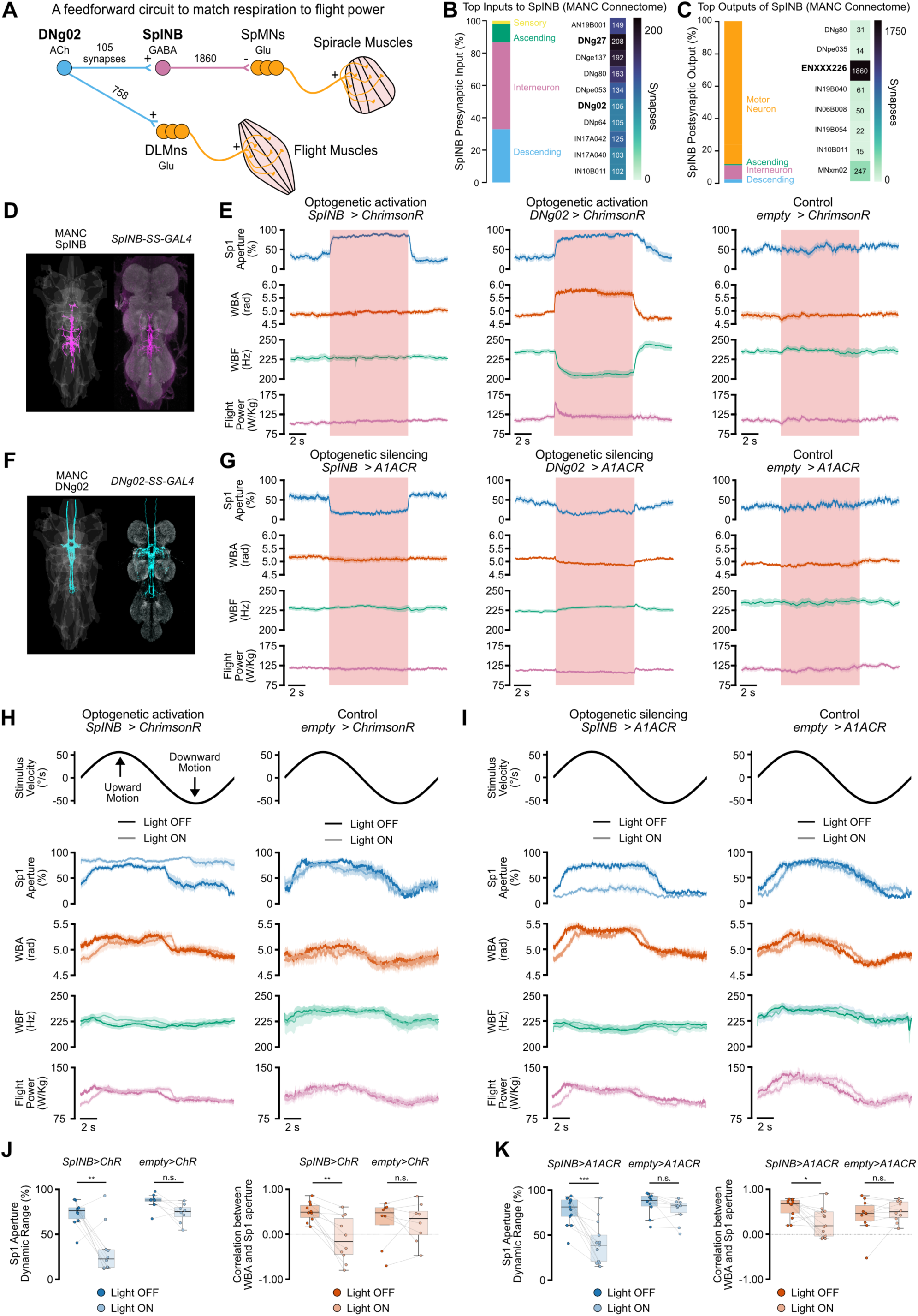
A flight control descending pathway opens spiracles, and SpINB is required to match spiracle aperture to flight power. **(A)** DNg02 connects to SpMNs via SpINB and makes direct excitatory synapses onto DLM motor neurons. **(B and C)** SpINB’s presynaptic input and output are dominated by intrinsic and descending neurons, respectively. **(D and F)** MANC skeletons and GAL4 expression patterns confirm SpINB and DNg02 identity in the VNC. **(E)** SpINB activation opens Sp1 without affecting WBA or WBF; DNg02 activation opens Sp1 while also increasing WBA and transiently flight power. n = 10 each. **(G)** SpINB silencing closes Sp1, indicating tonic SpINB activity during flight; DNg02 silencing closes Sp1 and reduces WBA and flight power. n = 11, 20, 11. **(H and I)** With SpINB activation, Sp1 aperture is clamped open and no longer tracks the optomotor stimulus, while wing kinematics continue to track normally; with SpINB silencing, aperture is clamped near closed with similarly attenuated visual modulation. Controls are unaffected in both cases. **(J and K)** SpINB activation and silencing each disrupt the normal correlation between WBA and Sp1 aperture and reduce the dynamic range of aperture during optomotor stimulation; controls are unaffected. Paired Wilcoxon signed-rank tests; *p < 0.05, **p < 0.01, ***p < 0.001.

We identified a Split-Gal4 driver that labeled the pair of SpINB interneurons and confirmed its expression (**Fig. 5D**). Optogenetic activation of SpINB opened spiracles in flying flies without changing wingbeat frequency or amplitude (**Fig. 5E, Movie S9**), consistent with its predicted GABAergic (inhibitory) output onto SpMNs. If descending flight commands act through this pathway, activating DNg02 itself should also open spiracles. Indeed, optogenetic activation of DNg02 simultaneously increased flight power and opened the spiracles in flying flies (**Fig. 5F-G, Fig. S4A, Movie S10**). DNg02 activation opened the spiracles even in immobilized, wingless flies that were not generating flight power (**Fig. S4B**). Because this opening occurs in the absence of flight, it cannot be explained by reafferent sensory feedback (e.g., falling O₂ or rising CO₂), but is more likely a feedforward copy of the descending motor command. This architecture allows the nervous system to predictively match spiracle aperture to the commanded level of motor exertion, rather than relying on slower interoceptive feedback.

If SpINB cells match spiracle opening to flight power, silencing these neurons during flight should close the spiracles and decouple spiracle aperture from flight motor output. Acute optogenetic silencing of SpINB cells closed the spiracles during flight, indicating that they are active and provide ongoing drive to open the spiracles during flight (**Fig. 5G, Movie S9**). Moreover, during the optomotor lift response (**Fig. 2B**), optogenetically silencing SpINB cells clamped Sp1 near its closed state and strongly attenuated its visual modulation, reducing both the dynamic range of spiracle aperture and its covariation with WBA, while wing kinematics were unaffected (**Fig. 5I, K**). Conversely, optogenetic activation of SpINB cells disrupted the normal relationship between spiracle aperture and WBA without altering wing kinematics (**Fig. 5H, J**). These results demonstrate that SpINB neurons couple the descending flight command to spiracle opening.

### A feedback circuit for metabolic regulation of spiracle size

We next sought to identify a neural circuit positioned to sense internal state and decrease spiracle opening during flight. Analysis of SpINB presynaptic inputs identified DNg27, a pair of glutamatergic descending neurons, as a candidate node linking flight, metabolism, and spiracle control (**Fig. 6A, B**). In the male CNS connectome, DNg27 receive strong input from interoceptive sensory neurons (ISNs) (*57*, *58*), which sense hemolymph osmolarity and adipokinetic hormone (AKH), an insect hormone that signals nutrient deprivation, analogous to glucagon in mammals. DNg27 cells also receive strong input from corazonin-expressing neurosecretory cells implicated in desiccation and metabolic stress (*59–61*). These inputs suggest that DNg27 is positioned to relay information about water and glucose homeostasis from the brain to the VNC. In the VNC, DNg27 cells synapse on neurons involved in spiracle or flight control, including SpINB, SpMNs and DLMns.

**Fig. 6.**
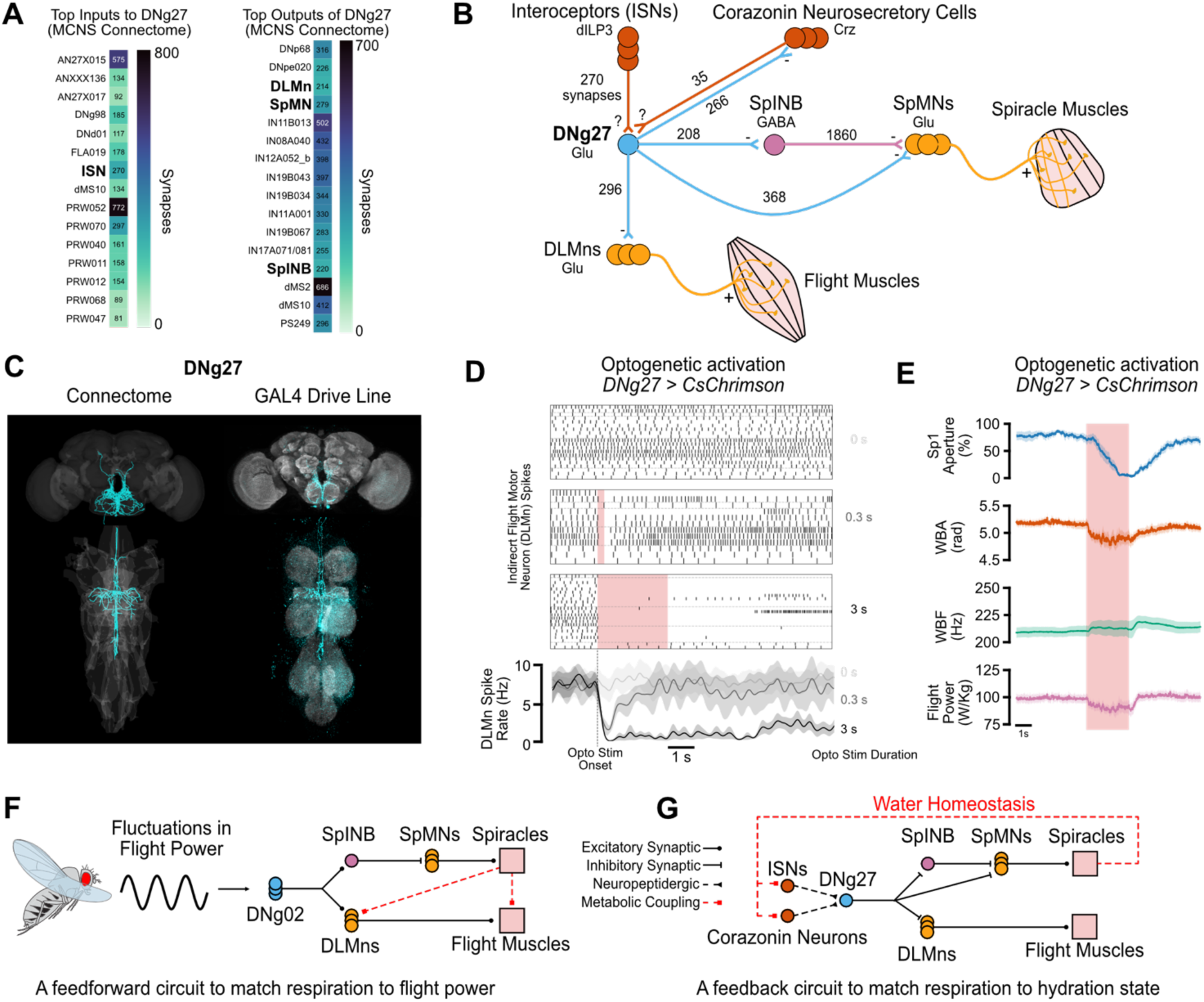
A candidate interoceptive pathway that closes spiracles and reduces flight power. **(A)** DNg27’s top inputs include interoceptive sensory neurons (ISNs), which respond to hemolymph osmolarity and AKH signaling; top outputs include DLMns, SpMNs, and SpINB. **(B)** Connectome-derived circuit schematic. Corazonin-expressing neurosecretory cells (l-NSC^Crz^), implicated in water homeostasis, also synapse onto DNg27. **(C)** DNg27 skeletons and GAL4 expression confirm its identity in brain and VNC. **(D)** DNg27 activation silences DLMn firing, consistent with direct glutamatergic inhibition. n = 5–6. **(E)** DNg27 activation closes Sp1 with slower kinematics and reduces wingbeat amplitude and flight power. n = 9. **(F)** Proposed feedforward module: visual lift commands via DNg02 drive both spiracle opening (via SpINB/SpMNs) and flight power (via DLMns), coupling aperture to instantaneous flight demand. **(G)** Proposed interoceptive feedback module: ISN and Corazonin cell input onto DNg27 drives spiracle closure while suppressing DLMn output, a candidate mechanism by which water status constrains respiratory water loss. Together, the two modules tune spiracle aperture to both metabolic demand and water balance.

We used a DNg27-GAL4 driver (**Fig. 6C**) to test if DNg27 neurons regulate flight or spiracle aperture. Activation of DNg27 silenced DLMns during flight, consistent with its predicted neurotransmitter (glutamate) identity and direct synaptic connections onto flight motor neurons (DLMns, **Fig. 6D**). Optogenetic stimulation of DNg27 frequently terminates flight altogether. However, in the flies that sustained flight after three seconds of DNg27 activation, the spiracles were closed, and wingbeat amplitude and flight power decreased (**Fig. 6E, Movie S11**). DNg27 neurons synapse directly onto SpMNs, but they also provide strong input to SpINB, whose inhibition is sufficient to close the spiracles (**Fig. 5G**).

Overall, these findings identify two neural pathways for coordinating respiration during flight: a descending feedforward command (DNg02) that predictively matches spiracle aperture to metabolic demand during flight (**Fig. 6F**), and a descending interoceptive feedback pathway (DNg27) that closes the spiracles and suppresses flight output during dehydration (**Fig. 6G**).

## Discussion

Using high-speed spiracle imaging, connectomics, optogenetics, and electrophysiology, we identify a compact circuit that controls respiration during flight in *Drosophila*. A descending pathway that commands flight power, DNg02, also opens the spiracles, matching respiration to locomotor demand. A separate interoceptive pathway, DNg27, can close the spiracles and suppress flight. Both converge on the same GABAergic interneuron, SpINB, which regulates the activity of the spiracle motor neurons (SpMNs) and is positioned to reduce gas exchange when water conservation takes priority over oxygen supply.

We used the fly VNC connectomes to identify the SpMNs and manipulated their activity directly in behaving flies. Activating SpMNs closed the spiracles and silencing them opened the spiracles, showing that *Drosophila* spiracles are controlled by closer muscles. This result is consistent with classic anatomical descriptions in other dipterans (*62*), but distinct from locusts and cockroaches, which possess separate opener and closer muscles (*63*, *64*). Because flies lack an opener muscle, we hypothesize that spiracle opening is passive: when the closer relaxes, the resilin-rich flaps that guard the orifice (**Fig. 1C**) act as an elastic spring that restores the open state.

High-speed imaging of individual spiracles revealed that thoracic aperture continuously tracks fluctuations in flight power, with a phase lag comparable to that of wingbeat amplitude. This confirms the central conclusion of earlier respirometry studies, which found that spiracle opening scales with metabolic demand during flight (*4*). It further demonstrates that this respiratory-to-locomotor matching is fast and continuous. Such coupling was first observed in dragonflies 60 years ago, where a central inhibitory reflex was proposed to open the spiracles at flight onset even in the absence of flight movements (*45*, *46*). However, the circuit basis of flight-matched spiracle control was unknown. We find that DNg02 neurons drive the wing motor system and, through SpINB, also controls spiracle aperture.

A defining feature of the circuit that matches spiracle opening to flight demand is that it is feedforward. DNg02 opens the spiracles even in immobilized flies that produce no flight power (**Fig. S4B**), indicating that opening reflects a copy of the descending command rather than a reaction to its metabolic consequences. This makes sense given the short timescale: tracheal and hemolymph gas stores buffer less than a second of flight metabolism (*43*), so a controller that waited for internal oxygen to fall would be too slow to keep pace with demand. The same constraint applies to any organ supplying oxygen to working muscle. The vertebrate cardiovascular and respiratory systems appear to have converged on analogous solutions: ventilation and cardiac output rise at the onset of exercise, driven by central commands from locomotor regions rather than by feedback from the muscles (*9*, *65*). That insects and vertebrates have the same predictive strategy, despite very different respiratory and circulatory architectures, suggests that feedforward control of metabolic support is a convergent feature of motor systems. A distinctive property of insects is that the tracheal system delivers oxygen directly to the tissues, so the spiracle is at once the site of gas exchange and the point of control, whereas vertebrates distribute these functions across the lungs, heart, and vasculature.

Closing the spiracles did not simply silence the flight motor program. When we activated SpMNs during flight, motor neuron firing fell only modestly while flight power dropped more steeply. This dissociation indicates that spiracle closure impairs flight largely through a peripheral failure of muscle force production, consistent with oxygen limitation of the flight muscles. Insect flight muscles sustain some of the highest known metabolic rates (*20*, *22*). Our measurements could not distinguish whether the decrease in muscle force production reflects oxygen depletion, accumulation of carbon dioxide, or both. In other insects, both gases influence spiracle control, and carbon dioxide set-points feature prominently in models of cyclic gas exchange (*66*). The bottleneck nonetheless lies downstream of the motor circuit, in the muscle’s capacity to convert spikes into force when gas exchange is restricted. The modest reduction in motor neuron firing during spiracle closure also hints at a feedback signal that reports the metabolic state of the flight muscles back to the nervous system. Interoceptive sensory neurons that monitor muscle metabolism would be well placed to provide this feedback signal, but no such neurons have yet been identified in the *Drosophila* flight muscles. Current connectomes are limited to the central nervous system and would not capture such peripheral sensors. A technique that can image whole peripheral tissues at cellular resolution, such as X-ray nanotomography (*67*), could survey the flight muscles for these candidate sensors.

Spiracle aperture is also modulated on a slower timescale by internal state. Following desiccation, flight performance declines and spiracle aperture is reduced (**Fig. 2E-G**). DNg27 provides a strong candidate pathway that links interoception and spiracle closing. Activating DNg27 closes the spiracles and suppresses flight, reproducing the direction of both effects seen after desiccation, and our connectome analysis together with prior work indicates that DNg27 receives input from interoceptive sensory neurons (ISNs) and corazonin neurons, which sense water and metabolic state (*58–61*). Both ISNs and corazonin neurons release neuropeptides, so determining the sign and function of their signaling to DNg27 will be important for understanding how the fly balances oxygen delivery against water conservation.

Our findings raise several new questions about insect respiration. The connectome points to highly synchronized control between left–right pairs and between the thoracic spiracles, but the distinct inputs to abdominal SpMNs, and the smaller difference between Sp1 and Sp2 inputs (**Fig. S2F, G**), suggest that different segments may be controlled separately. This organization could contribute to directional airflow, as proposed in blowflies (*16*). The spiracles also close slowly after flight (**Fig. 1F**), suggesting that steady-state aperture is shaped by internal gas state in addition to the feedforward command. This slow closing could reflect a gradually discharged oxygen debt. The spiracles flutter continuously during flight, yet the coupling between spiracle aperture and any single flight variable is weak on short timescales. How the nervous system finely controls spiracle aperture to admit enough oxygen while minimizing water loss, and what generates the flutter, remain unknown.

More broadly, by identifying and gaining genetic access to the SpMNs and their upstream neurons, this work provides a new entry point to understand the neural control of insect respiration. It is now possible to ask what additional signals drive changes in spiracle aperture and how restricting respiration affects other behaviors such as walking, sleep, and feeding. Because the same driver lines may label SpMNs in other drosophilids adapted to arid or humid niches, they could also allow respiratory circuits to be compared across species and reveal how spiracle control has evolved with ecology. As climate change increases the temperature and desiccation stress faced by insects, understanding how the nervous system balances oxygen uptake against water loss may help explain why some species tolerate aridity better than others (*40*).

## Acknowledgements

We thank members of the Tuthill Lab for technical assistance and feedback on the manuscript. We thank the members of the 2024 Neural Systems and Behavior (NS&B) Course at MBL, where spiracle imaging pilot experiments were initially conducted. We thank Tom Daniel and Michael Reiser for helpful discussions and generous loaning of equipment. We thank Bing Brunton, Sama Ahmed, and Nathan Baertsch for comments on the manuscript. We thank Jan-Marino Ramirez, Jeffrey Riffell, John Carlson, Aguan Wei, and Meet Zandawala for constructive feedback during the course of these experiments. We thank Tsunaki Asano for generously sharing the resilin-GAL4 and UAS-resilin fly lines.

## Funding

This work was supported by the Marine Biological Laboratory (MBL), the Neural Systems & Behavior Course (NS&B), NIH grant K99NS146574 to Y.L., and NIH grant R01NS128785, a Searle Scholar Award, a Klingenstein-Simons Fellowship, a Pew Biomedical Scholar Award, a McKnight Scholar Award, a Sloan Research Fellowship, the New York Stem Cell Foundation, and a UW Innovation Award to J.C.T. J.C.T is a New York Stem Cell Foundation – Robertson Investigator.

## Author Contributions

Y.L. and J.C.T. conceived the study and designed the experiments. Y.L. performed the high-speed spiracle imaging, tethered-flight behavioral experiments, optogenetics experiments, electrophysiology experiments, connectomic analyses, and analyzed all the data. A.S. performed the immunohistochemistry and confocal imaging. J.I. performed the scanning electron microscopy. G.K. and Y.Y. contributed to piloting the tethered-flight and optogenetic experiments. F.v.B. designed and built the free-flight arena, developed the trajectory-tracking and flight-analysis pipeline, and supervised the free-flight experiments and their analysis. Y.L. and J.C.T. wrote the manuscript with input from all authors. J.C.T. supervised the project and acquired funding. All authors read and approved the final manuscript.

## Declaration of Interests

The authors declare no competing interests.

## Data and code availability

Data is available on Dryad (https://doi.org/10.5061/dryad.zkh1893s4). Code for analyzing high-speed spiracle videos, EMG data, or connectomics data is located on GitHub (https://github.com/camellyc).

## Supplementary Videos

**Movie S1.** Continuous imaging of Sp2 during a brief (∼1 s) flight bout, recorded at 210 fps so the flapping of the wing is visible in the video and obscures the spiracle. Sp2 opens at flight onset, remains open throughout the bout, and closes after flight cessation. Note that this video is not motion corrected and for illustration purposes only (i.e., not used for spiracle aperture quantification).

**Movie S2.** Continuous imaging of Sp1 during a brief (∼3 s) flight bout, imaged at a constant 1,000 fps so the flapping of the wing is visible in the video and partly obscures the spiracle. Sp1 opens at flight onset, remains open throughout the bout, and closes after flight cessation. For quantitative analysis, frames in which the spiracle is obscured were identified and excluded *post hoc* based on average frame intensity. Corresponds to Fig. 1F.

**Movie S3.** Sp1 responding to an oscillating visual stimulus; stimulus velocity and Sp1 aperture are plotted below. Frame acquisition was triggered on each wingstroke, so the wing did not obscure the spiracle (same for subsequent videos). Movie is replayed at 200 fps, so playback is roughly realtime.

**Movie S4.** Sp2 responding to an oscillating visual stimulus.

**Movie S5.** (Left) Fluttering Sp1 during flight (30 minutes after tethering). Corresponds to Fig. 2E. (Right) Fluttering Sp1 during flight after 30 minutes desiccation (60 minutes after tethering). Corresponds to Fig. 2F.

**Movie S6.** GluRIIB-GAL4 labelling showing that a single SpMN innervates each spiracle closer muscle. **(A)** Sp1, **(B)** Sp2, **(C)** an abdominal spiracle.

**Movie S7.** Optogenetic activation of SpMNs (SpMN-SS1 > ChrimsonR) closes **(A)** the Sp2 and **(B)** an abdominal spiracle.

**Movie S8.** Optogenetic activation (**A**, SpMN-SS1 > ChrimsonR) and silencing (**B**, SpMN-SS1 > A1ACR) during flight. **Movie S9.** Optogenetic activation (**A**, SpINB-SS > ChrimsonR) and silencing (**B**, SpINB-SS > A1ACR) during flight. **Movie S10.** Optogenetic activation (**A**, DNg02-SS > ChrimsonR) and silencing (**B**, DNg02-SS > A1ACR) during flight. **Movie S11.** Optogenetic activation in DNg27 > CsChrimson flies.

## Materials and Methods

### Experimental animals

*Drosophila melanogaster* were reared on standard cornmeal-glucose medium (Archon Scientific) at 25 °C under a 12 h:12 h light:dark cycle. All experiments used mated females, 2-10 days post-eclosion. For optogenetic experiments, flies were transferred 1-3 days before experiment to vials containing cornmeal-glucose medium supplemented with 80 µL of 35 mM all-*trans*-retinal in 95% ethanol (Santa Cruz Biotechnology) 1-3 day before the experiment, and kept in the dark until use.

### Immunohistochemistry

For immunohistochemistry of the fly CNS, female flies were cold anaesthetized on ice, then dissected in cold PBS. After 55 min fixation of samples in 2% PFA/PBS, samples were washed 4 times for 15 mins in 0.3% PBST at RT, blocked in 5% normal goat serum in 0.2% PBST for at least 1.5 hrs, and incubated with primary antibodies diluted in 0.2% PBST. Samples were washed again 4 times for 15 mins in 0.3% PBST at RT, and incubated with secondary antibodies for 2 days in darkness. Samples were then washed in 0.2% PBST, and then mounted on a slide with Vectashield (Vector Laboratories). We acquired z-stacks of each VNC on a confocal microscope (Olympus FV1000). We aligned the morphology of the VNC to a female VNC template in ImageJ with the Computational Morphometry Toolkit plugin (CMTK32; http://nitrc.org/projects/cmtk).

For imaging GFP expression in spiracles, we fixed intact flies (4% paraformaldehyde in PBS) for 30 minutes. We washed them 3x in PBS. Next, we hemisected flies by transferring them one by one to a small drop of Tissue-Tek O.C.T., freezing them on a glass slide on dry ice for 15 seconds, and slicing them along the anterior-posterior axis with a fine razor blade. We immediately transferred them to a well containing PBS for one minute, until the O.C.T. melted and dissolved. The tissue was then washed 3 times in PBS containing 0.2% Triton-X (PBST) over one hour. To label muscles, tissue was incubated overnight in Alexa Fluor 647-nm Phalloidin in PBST. Tissue was washed three times with PBST over three hours, once in PBS for one hour, and mounted in Vectashield on a slide with four layers of 3M double-stick tape for spacers. Preps were imaged on an Olympus FV1000 confocal.

### Scanning electron microscopy

Whole flies were cold anesthetized, dipped briefly in PBS-T 0.1% and rinsed in PBS to promote sinking in the processing solutions. Specimens were incubated over night at 4C in fix (2% formaldehyde, 2.5% glutaraldehyde in 0.1 M cacodylate buffer). To capture the spiracles in their closed configuration, *SpMN-SS1>ChrimsonR* flies were used, and red light was applied during the first 15 min of the fixation process. After fixation, samples were rinsed with 0.1 M cacodylate buffer, and post-fixed in 1% OsO4 in 0.1 M cacodylate buffer for 1 hour. Samples were then washed in distilled water and gradually dehydrated by sequential 5-min incubations in ethanol at 35%, 50%, 70%, 95%, 100% and 100%. Specimens were then subjected to critical point drying (Denton CDP-1, Denton Vacuum Inc.) and mounted in a 12 mm diameter pin stub (Ted Pella Inc.) using double sided carbon tab. Special care was taken to orient the flies on their lateral side, exposing the spiracles. Samples were then sputter-coated with 20 nm gold deposition. Scanning electron micrographs of the spiracles were acquired using an Apreo S microscope (ThermoFisher).

### Tethered flight behavior

Flies were cold-anaesthetized at 2.5 °C and tethered dorsally on the thorax with a 0.005-inch tungsten pin (A-M Systems, #556892) using UV-curable glue (KOA 300, Kemxert; thicker Bondic UV glue for EMG experiments). Tethered flies were mounted in the center of a cylindrical LED arena (*68*) (height 128 mm, diameter ∼125 mm) within a dark, climate-controlled chamber held at 24-25 °C and 55-60% relative humidity. Flies were habituated in the arena for ≥30 min before recording unless otherwise noted. Each experimental trial lasted 60 – 180s, depending on the experiment. At the start of each trial, flies were allowed to fixate on a vertical dark bar displayed on the LED panels. At the end of each spiracle imaging trial, a brief pulse of 100% CO₂ was delivered to fully open the spiracles; in *SpMN-SS1>ChrimsonR* flies, a 0.5 s optogenetic pulse was additionally used to confirm complete spiracle closure.

Wingbeat kinematics were measured with an optical wingbeat analyzer as described elsewhere (*69*). Briefly, an infrared LED above the fly projected the wing shadows onto a pair of photodiodes, generating an analog waveform (“hutchens”) whose frequency and amplitude were extracted in real time as wingbeat frequency (WBF) and per-wing wingbeat amplitude (WBA_L_, WBA_R_). The left-minus-right WBA signal was fed back to the LED arena to control the velocity of the closed-loop vertical bar; the left-plus-right WBA signal was recorded as total WBA. WBA and WBF signals detected by the wingbeat analyzer were then smoothed with a 5-ms moving-average window. For experiments requiring quantification of flight power and thus WBA in radians, WBA was extracted and calculated from video. A Basler camera fitted with an InfiniStix lens recorded a top view of the fly perpendicular to the wingstroke plane at 100 fps, and frames were analyzed post-hoc with a modified version of Kinefly (*70*, *71*) to extract WBA in radians. Flight power was computed following Gordon et al. (2006) as the sum of profile and induced power, *P_mech_* = *P_pro_ + P_ind_*, with *P_pro_* = *C_pro_Φ³n³* and *P_ind_* = *C_ind_Φ^-1/2^*, where Φ is mean wingbeat amplitude (rad), n is wingbeat frequency (Hz), *C_pro_* = 1.44 × 10⁻⁷ m² and *C_ind_* = 10.5 m²s⁻³, and the flight muscle is assumed to account for 30% of total body mass.

For the 30 vs 60-min comparison (Fig. 2E-G), flies were imaged after the standard 30-min habituation, then transferred to a vial containing Drierite desiccant for 30 min and re-imaged. For the optomotor protocol, a closed-loop vertical dark bar was superimposed on open-loop horizontal dark stripes; the horizontal stripes oscillated vertically with sinusoidal velocity (period 20 s, peak velocity 56°·s⁻¹). For optogenetic experiments, a fiber-coupled LED (M625F2, Thorlabs) driven by a current driver (LEDD1B, Thorlabs) delivered light through a 200 µm-diameter optic fiber (M92L01, Thorlabs) and a custom-made collimator (*72*) (MAP10100100-A, Thorlabs) positioned over the thorax, with care taken to minimize illumination of the head. Stimuli were delivered as 200 Hz pulse trains (3 ms pulse width) at ∼150 µW·mm⁻² over a ∼1 × 1 mm spot covering the entire thorax, timed by TTL pulses generated by a custom script through a NI PCIe-6321 board.

### Spiracle imaging

Spiracles were imaged with a Phantom KT-810 high-speed camera fitted with an InfiniTube FM-100 tube lens, a Nikon L-Plan ELWD 50×/0.45 objective, and a Schott RG-780 long-pass filter. The fly was illuminated with an M850F3 fiber-coupled LED (Thorlabs) focused with a custom collimator (*72*).

For immobilized fly imaging, legs were amputated at the coxa and wings at the wing base, and the thorax and abdomen were immobilized with UV-curable glue. Images were captured at 512 × 512 pixels over a ∼200 × 200 µm field of view with a 150 µs exposure.

During flight, spiracle frames were similarly acquired at 512×512 pixels over a ∼100×100 µm field of view with a 150 µs exposure. Cameras were either internally triggered at 1000 fps to capture pre-flight, in-flight and post-flight spiracle dynamics (Fig. 1F), or externally triggered at wingbeat frequency using a custom-built converter that converted the per-wingbeat signal into a TTL pulse train at the same frequency. Trigger signals were recorded to align video frames with wingbeat frequency and amplitude. The forelegs were removed to prevent them from obstructing the optical path during imaging.

Sp1 aperture remained largely stable within each wingstroke cycle, and we observed no obvious wingstroke-phase-locked modulation, although the phase during which the wing occludes the spiracle completely could not be assessed. Care was taken to ensure robust triggering on each wingbeat. Trigger signals were digitized on a NI PCIe-6321 board and aligned post-hoc to WBA and WBF. Sp1 aperture was quantified as the mean pixel intensity within an ROI over the Sp1 aperture, divided by the mean intensity of a ROI in the adjacent cuticle to correct for fluctuations in the intensity of illumination light. Per-trial, the 1st and 99th percentile intensities were defined as 0% and 100% aperture, respectively, and all values were linearly rescaled within those bounds. Because the spiracle is a three-dimensional structure whose flaps rarely lie parallel to the imaging plane, the focal plane was chosen such that the edges of both flaps were clearly resolved at ∼50% Sp1 aperture.

Spiracle aperture typically shows substantial modulation ∼30 min after tethering. Because every trial is rescaled to the 1st and 99th percentiles of mean intensity within the spiracle ROI, good calibration depends on the fly fully closing its spiracle at some point during the trial. In *SpMN-SS1 > ChrimsonR* flies, complete closure can be driven reliably with an optogenetic pulse, so these recordings are better calibrated than other manipulations. In all other flies, spiracle closure was verified by the experimenter reviewing the full video of each trial.

For spiracle imaging with internal, fixed frame rate recording in Fig. 1F, videos were recorded at 1,000 fps. Videos were recorded and processed offline to extract per-frame spiracle aperture measurements. Raw video was shown in Supplementary Video 1. To extract spiracle aperture data and exclude frames in which the spiracle was transiently occluded by the wing, two ROIs (Sp1 and reference) were monitored throughout each trial. For each ROI, the mean pixel intensity at each frame was compared with its 51-frame centered rolling median, and any frame in which the wing blocks the spiracle (relative deviation of exceeded 5%) in either ROI was flagged and removed. The aperture values retained after this filtering step were then min-max rescaled to [0, 1] similar to externally triggered videos.

For the *DNg27 > CsChrimson* experiments in Fig. 6E, a 3 s stimulus stopped flight in most flies. Because the camera was triggered on wingbeats, no frames were captured once flight ceased, which makes spiracle aperture impossible to record. We therefore included only trials in which the 3 s stimulus did not fully stop flight (∼20% of trials).

### Oscillating visual stimulus analysis

To quantify how flight and spiracle signals tracked the oscillating optomotor stimulus, each response signal (wingbeat amplitude, wingbeat frequency, inferred flight power, and Sp1 aperture) was compared to the sinusoidal stimulus velocity (period 20 s, f = 0.05 Hz), modeled as *s(t) = sin(2πft)* with zero-crossings at the onset of each of the three 20 s cycles within a trial. For each cycle, we estimated the amplitude and phase of the response at the stimulus frequency by lock-in projection, computing *a = 2⟨y(t)·cos(2πft)⟩ and b = 2⟨y(t)·sin(2πft)⟩* over the finite samples of the cycle, equivalent to a least-squares fit of *y(t) = R·sin(2πft + φ)*, with response amplitude *R = √(a² + b²)* and phase *φ =* atan2*(a, b)*. Response delay was computed as *τ = −φ/(2πf)*, with positive τ indicating that the response lagged the stimulus.

### Free flight experiments

Approximately 20 female flies were maintained on all-trans-retinal food and then starved for ∼8 h before the experiment. The flies were released into a wind tunnel with no airflow, and data were collected overnight; the flies had no access to food or water during the trial. The wind tunnel was similar in operation to one described previously(*73*), but with a slightly larger geometry (0.6 x 0.6 x 1.8 m working section), different cameras, and modified illumination, as described in more detail below.

The floor and walls of the wind tunnel are equipped with a high contrast backprojection film (spyeblack screen, spye.co). To provide the flies with a visual environment to navigate in, a black and white checkerboard was projected onto the floor of the wind tunnel from below (VA-LT002 4K UST Laser Projector, VAVA). The two side walls were back-lit with blue LEDs (ColorSpace™ RGB (blue channel only), Waveform Lighting). Both the walls and floor were back lit with 850 nm infrared LEDs to provide contrast for tracking (850 nm IR LED strip, Waveform Lighting). Two daylight (4500 K) white light strips (Centric Daylight, Waveform Lighting) with a diffuser were placed along the top edges of the tunnel to provide flies with sufficient long wavelength light for photoreceptor reisomerization. Optogenetic stimuli were controlled with a row of RGB LEDs (Triple Output High Power RGB LED (M018001MA3LZ), SparkFun) on each long side of the tunnel. To reduce the effect of a visual startle response, these LEDs were nominally set to a low level of blue illumination (10% intensity), and Chrimson channels were activated by turning on the red channel to 100% intensity. The red light irradiance at the center of the tunnel was 20.39 µW / mm^2^.

Flies were tracked in three dimensions at 100 fps using a real-time tracking system (*74*) with 20 Basler Ace2 cameras (a2a 1920-160um, Basler) equipped with Tamron lenses (Tamron 4-13 mm F/1.5 IR) and IR-pass filters (LEE Filters 3 x 3” Infrared (87C) Polyester Filter, bhphoto). A 0.2 × 0.2 × 0.6 m trigger zone was defined at the center of the chamber. Upon entry into this zone, the red channel of the RGB LEDs was delivered throughout the chamber on a randomly selected half of events, with the remaining half serving as no-light sham trials. The trigger code was set up such that there were always at least 10 seconds between triggers. For each trigger event, 0.2 s of pre-stimulus and at least 6 s, and up to 8 s of post-stimulus tracking were recorded, and the mean Z (vertical) position across events was computed using a custom Python script. Because the software could not maintain individual identities over the span of the experiment, each trajectory was treated as an independent sample.

### Extracellular electrophysiological recording of DLMn

Extracellular recording of the dorsal longitudinal flight motor neurons (DLMns) was performed as described elsewhere (*75*). A sharpened tungsten electrode was inserted dorsally into the dorsal longitudinal muscle of a tethered fly. No attempt was made to distinguish individual DLM fibers, as the DLMns are electrically coupled thus having the same firing rate during flight (*75*). A reference electrode of the same type was inserted between adjacent abdominal tergites. Signals were amplified 100× with an A-M Systems Model 1800 microelectrode AC amplifier and digitized at 20 kHz on the NI PCIe-6321 board. Spikes were detected with a custom python script. When more than one unit was detected, a single unit was isolated by thresholding on the spike-amplitude histogram.

### Connectomics

Connectivity was queried from the Male Adult Nerve Cord (MANC, v1.2.3) and Male Central Nervous System (MCNS v0.9) connectomes via neuPrint. Connections weaker than three synapses were excluded from analyses. For bilateral comparisons, left and right partners were matched by instance label (e.g. ENXXX226_T1_L with ENXXX226_T1_R). Where multiple neurons shared an instance, all pairwise L-R cosine similarities were calculated and plotted.

### Quantification and statistical analysis

All experiments were conducted or analyzed with custom scripts written in either Matlab or Python. Sample sizes refer to the number of individual flies tested; when multiple trials were performed in the same fly, the values were averaged and treated as a single value for that fly. Variance across individuals is represented by either a mean ± standard deviation trace, or by a box plot. Box plots follow the standard Tukey convention: the central line shows the median, the box spans the interquartile range (Q1 to Q3), and the whiskers extend to the most extreme observation within 1.5 × IQR of the nearest box edge. Individual data points are overlaid as jittered scatter. For paired comparisons, thin gray lines connect the two values belonging to the same fly across conditions.

### Use of AI tools

Claude (Anthropic) was used to assist in writing and debugging analysis code, and in editing manuscript text for clarity and consistency. The authors reviewed and verified all code, analyses, and text.

## Supplementary Materials

**Fig. S1.**
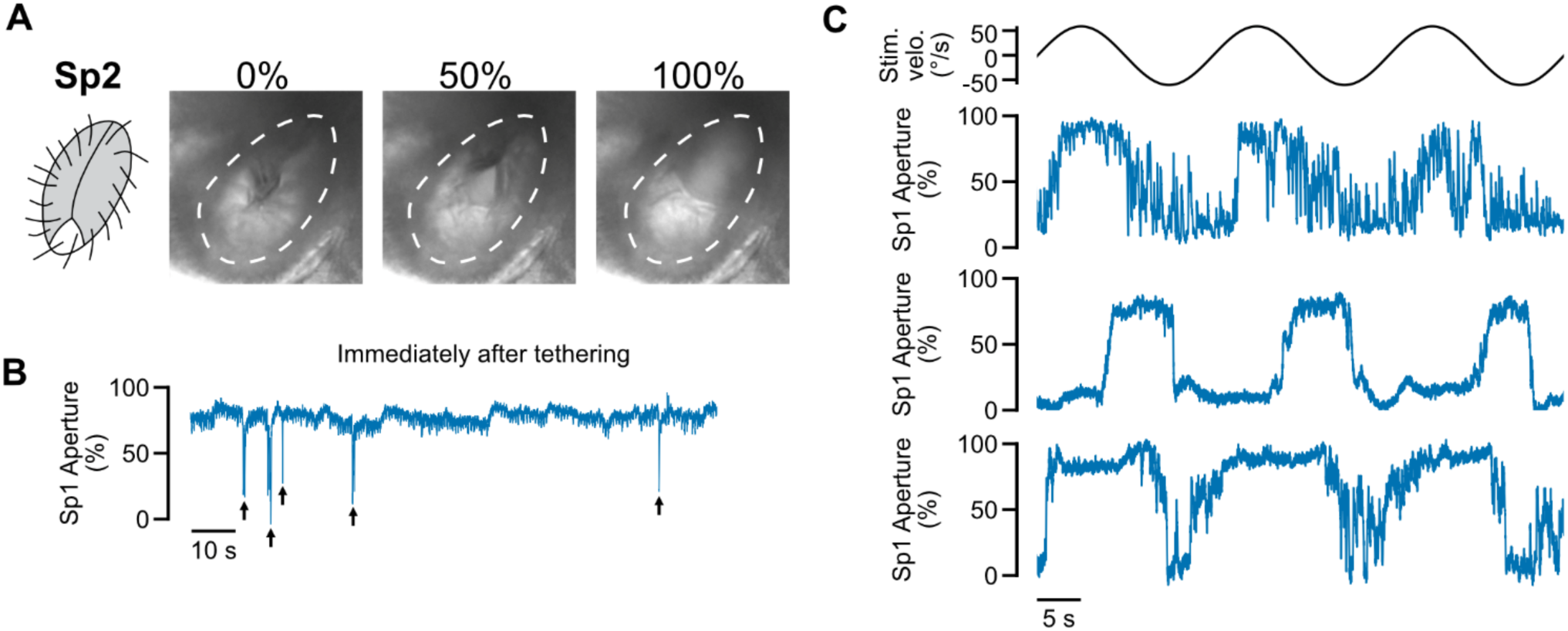
**(A)** Representative frames of Sp2 at 0%, 50%, and 100% aperture. **(B)** Representative Sp1 aperture recording from a single fly immediately after tethering. Sp1 is widely open and closes only briefly and occasionally (arrows). **(C)** Representative time series during optomotor stimulation in three individual flies; stimulus velocity is shown at top. Sp1 aperture generally follows the visual stimulus, but the depth of modulation and the phase delay vary across flies.

**Fig. S2.**
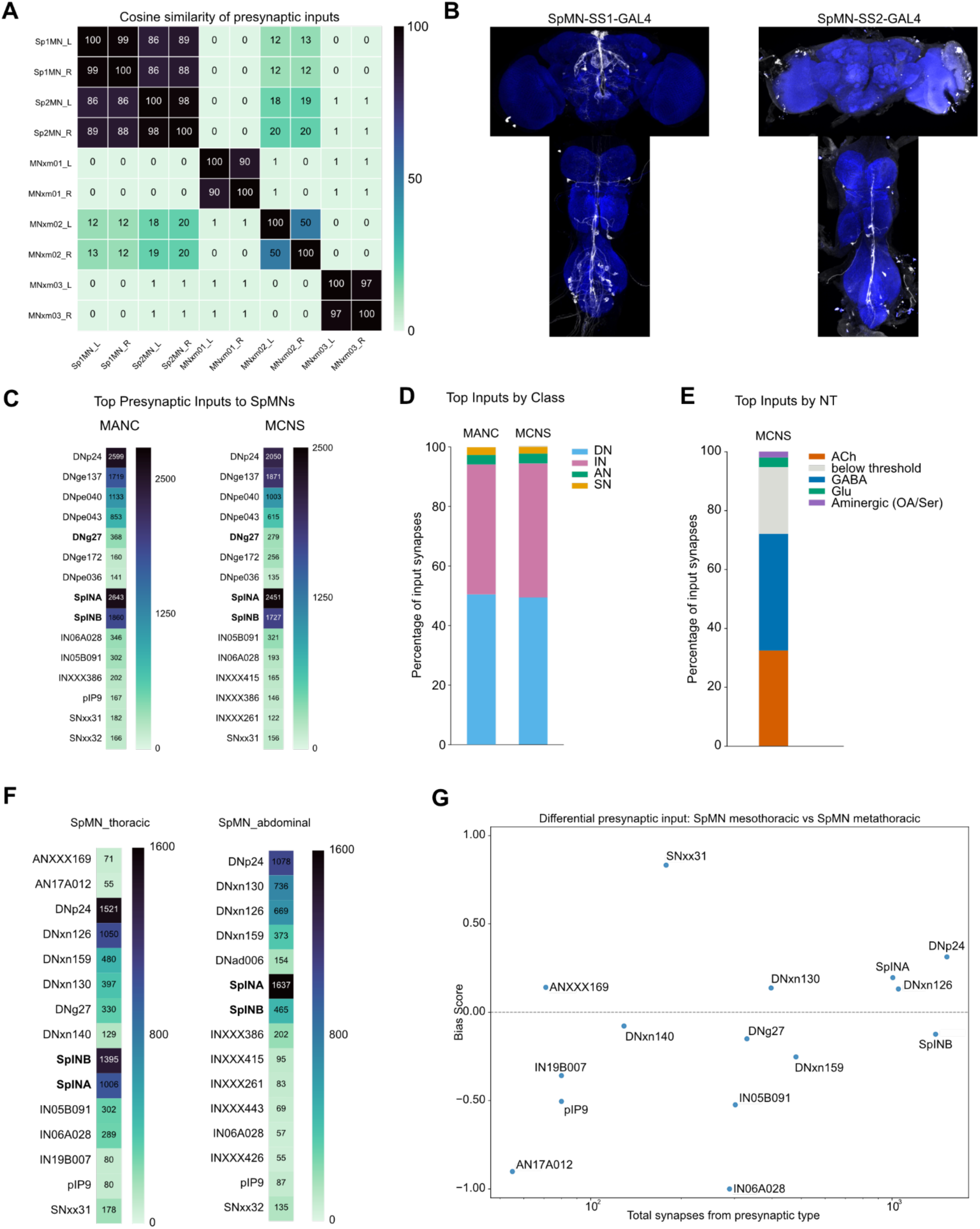
**(A)** Cosine similarity matrix of all unannotated efferent neurons projecting through the wing or haltere nerves. The two thoracic SpMN candidates (SpMN1, SpMN2) are highly similar to each other and distinct from MNxm01, MNxm02, and MNxm03, which have low similarity to one another. **(B)** Confocal expression patterns of the SpMN-SS1-GAL4 and SpMN-SS2-GAL4 driver lines. **(C)** Top 15 presynaptic inputs to SpMNs in the MANC and MCNS connectomes. **(D)** Composition of SpMN presynaptic input by neuron class in MANC and MCNS. **(E)** Composition of SpMN presynaptic input by predicted neurotransmitter in MCNS. **(F)** Top 15 presynaptic inputs to thoracic or abdominal SpMNs. **(G)** Bias score for presynaptic inputs to the mesothoracic (Sp1) versus metathoracic (Sp2) SpMN.

**Fig. S3.**
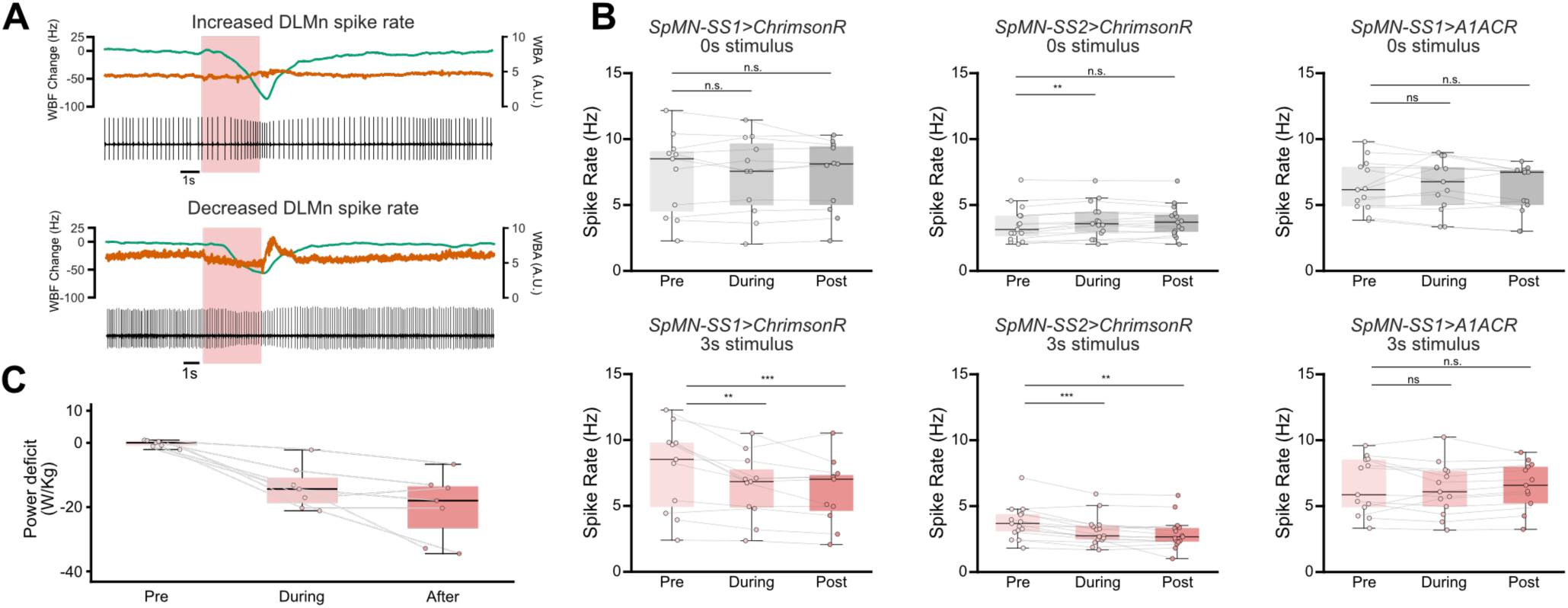
**(A)** Wingbeat frequency and amplitude can decline alongside either an increase or a decrease in DLMn spike rate during closure, indicating uncoupling of motor neuron firing from aerodynamic output. **(B)** DLMn spike rate is significantly reduced during and after closure for both SpMN-SS1 and SpMN-SS2 activation, but unchanged with silencing. n = 11 for *SpMN-SS1>ChrimsonR*, n = 15 for *SpMN-SS2 > ChrimsonR*, n = 13 for *SpMN-SS1 > A1ACR*. **(C)** Inferred flight power falls below the resting power–spike-rate baseline during and after closure, indicating reduced flight-muscle force output downstream of motor neuron firing. n = 7; paired Wilcoxon, *p < 0.05. **p < 0.01, ***p < 0.001; n.s., not significant.

**Fig. S4.**
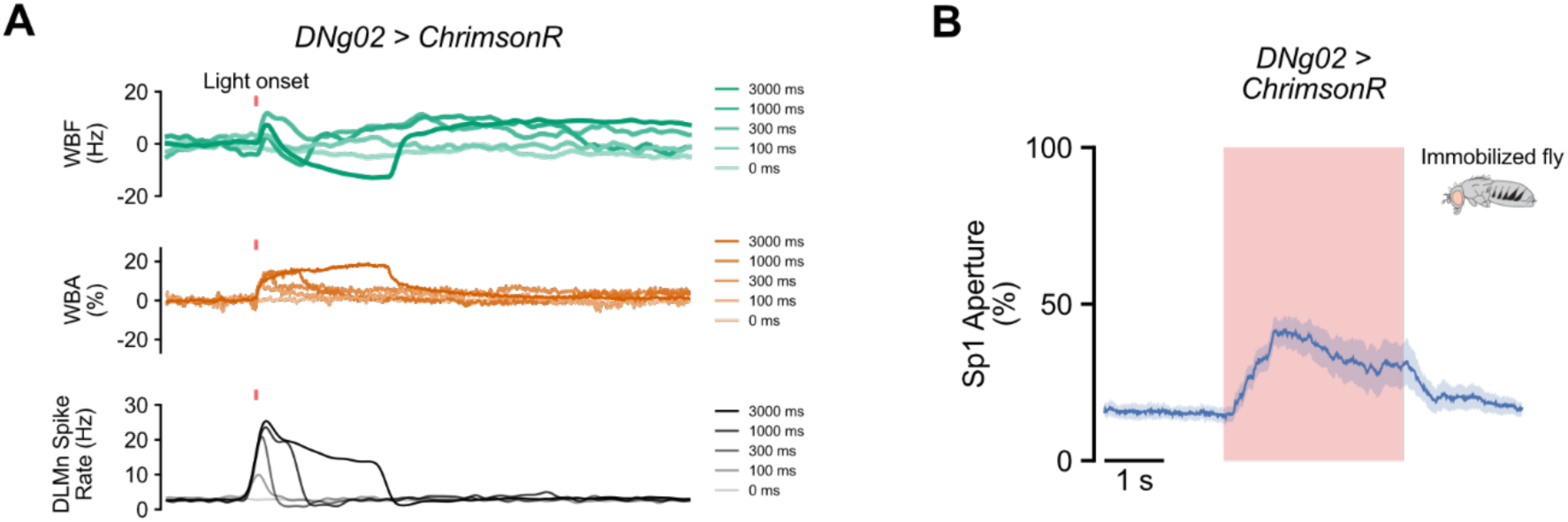
**(A)** Extracellular DLMn recording during DNg02 > ChrimsonR activation. DNg02 activation sharply increases DLMn spike rate, consistent with DNg02 being cholinergic and presynaptic to DLMns. **(B)** DNg02 > ChrimsonR activation in an immobilized fly. Sp1 opens in response to the optogenetic stimulus even in immobilized, non-flying flies. n = 10.

**table S1.** Key resources.

| Reagent or resource | Source | Identifier |
| --- | --- | --- |
| *Chemicals, peptides, and recombinant proteins* |  |  |
| Glucose Food in Narrow Vials | Archon Scientific |  |
| All-trans-retinal | Santa Cruz Biotechnology | Cat#SC-221196 |
| Paraformaldehyde | Fisher Scientific | Cat#15710-S; CAS: 30525-89-4 |
| Normal goat serum | Fischer Scientific | Cat#50197Z |
| Triton X-100 | FISHER SCIENTIFIC | Cat# AAA16046AE |
| Vectashield | Vector Laboratories | Cat#H-1000 |
| Tissue-Tek® O.C.T. Compound | Sakura Finetek USA | Cat#4583 |
| UV-curable glue | Kemxert | Cat# KOA 300 |
| UV-curable glue | Bondic |  |
| **Antibodies** |  |  |
| Chicken anti-GFP | Invitrogen | Cat# PA5-143569 |
| Mouse anti-brp (nc82) | Developmental Studies Hybridoma Bank | RRID: AB_2314866 |
| Goat anti-chicken Alexa 488 | Invitrogen | Cat # A-11039 |
| Goat anti-mouse Alexa 633 | Invitrogen | Cat # A-11001 |
| Alexa Fluor Phalloidin Alexa Fluor 647 (1:50) | Thermofisher | Cat # A22287 |
| **Software and algorithms** |  |  |
| ImageJ | Schneider et al | https://imagej.nih.gov/ij/ |
| CAVEclient | Dorkenwald et al | https://github.com/seung-lab/CAVEclient |
| MATLAB | MathWorks | https://www.mathworks.com/products/matlab.html |
| Python | Python Software Foundation | https://www.python.org/ |
| SciPy |  | https://scipy.org/ |
| CMTK (Computational Morphometry Toolkit) | Rohlfing & Maurer | http://nitrc.org/projects/cmtk |
| Kinefly | Suver et al | https://github.com/ssafarik/Kinefly |
| Benifly | Benjamin Cellini | https://github.com/bmslpsu/Benifly |
| **Deposited data** |  |  |
| Connectome data (MANC) | FlyEM - Janelia Research Campus | https://flyem.janelia.org/MANC |
| Connectome data (MCNS) | FlyEM - Janelia Research Campus | https://male-cns.janelia.org/ |

**table S2.** Fly stocks. Split-Gal4 hemidrivers, effectors, and other stocks used in this study.

| Stock genotype | Source or BDSC # | Notes |
| --- | --- | --- |
| w[1118]; P{y[+t7.7] w[+mC]=VT029591-p65.AD}attP40 | 72204 | Split half for SpMN-SS1 and SpMN-SS2 |
| w[1118]; P{y[+t7.7] w[+mC]=VT000355-p65.AD}attP40; P{y[+t7.7] w[+mC]=R47A08-GAL4.DBD}attP2 | 75868 | Split half for SpMN-SS1 |
| w[1118]; P{y[+t7.7] w[+mC]=R20F02-GAL4.DBD}attP2 | 69853 | Split half for SpMN-SS2 |
| w[1118]; P{y[+t7.7] w[+mC]=p65.AD.Uw}attP40; P{y[+t7.7] w[+mC]=GAL4.DBD.Uw}attP2 | 79603 | empty-SS-GAL4 control |
| w[1118]; P{y[+t7.7] w[+mC]=R42B02-p65.AD}attP40; P{y[+t7.7] w[+mC]=VT039465-GAL4.DBD}attP2 | 88990 | SS01562, DN<sub>g</sub>02-SS-GAL4, Namiki et al., 2018 |
| w[1118]; P{y[+t7.7] w[+mC]=VT037550-p65.AD}attP40 | 601342 | Split half for SpINB-SS-GAL4 |
| w[1118]; P{y[+t7.7] w[+mC]=VT017191-GAL4.DBD}attP2 | 73835 | Split half for SpINB-SS-GAL4 |
| w[1118]; P{y[+t7.7] w[+mC]=GMR13D04-GAL4}attP2 | 48559 | DN<sub>g</sub>27-GAL4 |
| w;+;10xUAS-ChrimsonR::mCherry | Janelia | ChrimsonR, outcrossed to Tuthill lab WT |
| w[1118]; P{y[+t7.7] w[+mC]=20XUAS-IVS-Syn21-A1ACR1-EYFP-p10}su(Hw)attP1 | 606146 | A1ACR, outcrossed to Tuthill lab WT |
| w;+;Resilin-GAL4 | Dr. Tsunaki Asano | resilin-GAL4 |
| w;+;UAS-Resilin-RA::mCherry | Dr. Tsunaki Asano | UAS-Resilin tagged with fluorescent protein. |

**table S3.**
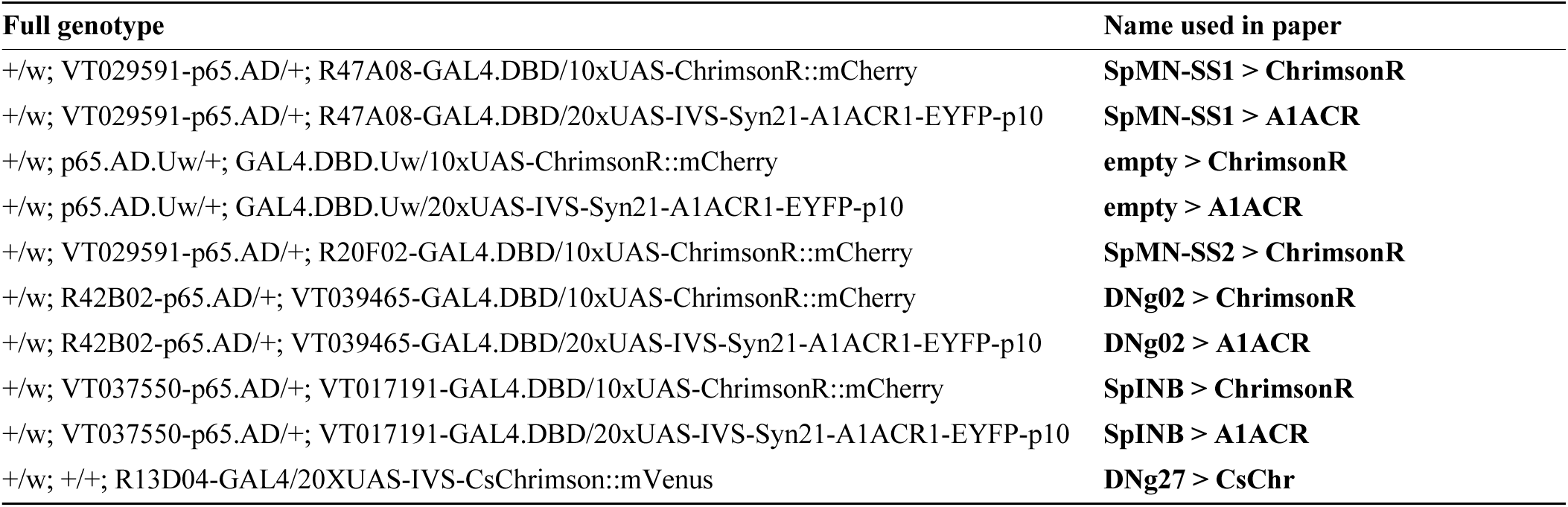
Driver line nomenclature. Full genotypes and the corresponding names used in the text and figures.

## Notes

https://doi.org/10.5061/dryad.zkh1893s4

### Competing Interest Statement

The authors have declared no competing interest.

